# Comparative hormonal regulatory pathway of the drought responses in relation to glutamate-mediated proline metabolism in *Brassica napus*

**DOI:** 10.1101/704726

**Authors:** Van Hien La, Bok-Rye Lee, Md. Tabibul Islam, Sang-Hyun Park, Dong-Won Bae, Tae-Hwan Kim

**Author notes:** These authors have contributed equally to this work (Van Hien La, Bok-Rye Lee). Correspondence: Tae-Hwan Kim. **Contact of other authors:** Van Hien La, Bok-Rye Lee, Md. Tabibul Islam, Sang-Hyun Park, Dong-Won Bae.

## Abstract

Proline metabolism influences metabolic and signaling pathway in regulating plant stress responses. This study aimed to characterize the physiological significance of glutamate (Glu)-mediated proline metabolism in the drought stress responses, focusing on the hormonal regulatory pathway. The responses of cytosolic Ca^2+^ signaling, proline metabolism and redox components to the exogenous application of Glu in well-watered or drought-stressed plants were interpreted in relation to endogenous hormone status and their signaling genes. Drought-enhanced abscisic acid (ABA) were concomitant with ROS and proline accumulation, accompanied by decreased NAD(P)H/NAD(P)^+^ and GSH/GSSG ratios. Exogenous Glu-feeding under drought resulted in an increase of salicylic acid (SA) with an antagonistic decrease of ABA. Glu-enhanced SA coincided with the highest expression of SA synthesis related gene *ICS1* and Ca^2+^-dependent protein kinase *CPK5*. SA-enhanced *CPK5* expression was closely associated with further enhancement of proline synthesis-related genes (*P5CS1, P5CS2*, and *P5CR*) expression. The Glu-activated proline synthesis was responsible for the reset of reducing potential with enhanced expression of redox regulating genes *TRXh5* and *GRXC9* in a SA-mediated *NPR1*- and/or *PR1*-dependent manner. These results clearly indicate that Glu-activated interplay between SA- and CPK5-signaling and Glu-enhanced proline synthesis are crucial in the amelioration of drought stress in *B. napus*.

**Highlight:**

- Drought-induced oxidative stress and symptom are developed by ABA-dependent manner
- Glu-application increases endogenous SA level with an antagonistic decrease of ABA
- Drought-induced proline accumulation was further enhanced by exogenous Glu-application
- Glu-enhanced proline synthesis accompanied with SA-mediated regulatory pathway
- Glu-enhanced SA-modulated proline metabolism is an integrated process of redox control

## Introduction

Prolonged water-deficit (e.g., drought) is considered a major climatic factor limiting plant growth and development. The decrease in water availability for transport-associated processes modifies intercellular metabolites concentration, followed by the disturbance of amino acid and carbohydrate metabolism (Kim *et al*., 2004; Lee *et al*., 2016). An accumulation of reactive oxygen species (ROS) and/or proline is observed as a common stress response (Lee *et al*., 2013; Rejeb *et al*., 2014). Indeed, rapid production of ROS (i.e., oxidative burst) is one of the earliest plant responses to stresses caused by a wide range of environmental stresses (Lee *et al*., 2009a) and pathogen infections (Finiti *et al*., 2014; Islam *et al*., 2017). Proline accumulation has been found to be also a primary stress responsive symptom resulting from dehydration in plant tissues such as drought conditions (Kim *et al*., 2004; Lee *et al*., 2009b), high salinity (Hong *et al*., 2000), or freezing temperature (Kaplan *et al*., 2007). The proline pool of plant cells depends on the rate-limiting steps in proline synthesis and degradation, which are catalyzed by Δ^1^-pyrroline-5-carboxylate synthease (P5CS) and proline dehydrogenase (ProDH) (Rejeb *et al*., 2014; 2015; La *et al*., 2019). Multifunctional roles of proline including in preventing oxidative damage, in stabilizing DNA, membranes and protein complex as well as in providing carbon and nitrogen source during stress have been well documented (Szabados and Savouré, 2010). Interestingly, proline metabolism has been reported to promote mitochondrial ROS production and enhance ROS in hypersensitive plants (Liang *et al*., 2013). Therefore, the modified proline metabolism by drought stress may further involves in drought stress tolerance by regulating intracellular redox potential (La *et al*., 2019), as well as energy transfer and reducing power (Szabados and Savouré, 2010; Rejeb *et al*., 2014), which are not yet fully understood.

Increasing evidences have shown that stress responsive ROS and/or proline metabolism are regulated by hormonal signaling pathways (Miura and Tada, 2014; Herrera-Vásquez *et al*., 2015; La *et al*., 2019). Among these, ABA-dependent signaling pathway has been more emphasized (Boudsocq and Sheen, 2013; Osakabe *et al*., 2014). Indeed, proline accumulation is partially regulated by an ABA-dependent signaling pathway in osmotic (Chung *et al*., 2008) and drought stress (La *et al*., 2019). Similarly, enhanced H_2_O_2_, as a ROS signaling from NADPH oxidase, stimulate ABA-induced proline accumulation (Verslues *et al*., 2007; La *et al*., 2019). Several studies have provided evidence for the ROS-mediated SA biosynthesis *via* Ca^2+^ signaling (Seyfferth and Tsuda, 2014; Herrera-Vásquez *et al*., 2015), as well as the proline-mediated biosynthesis of SA *via* NDR1-dependent signaling (Chen *et al*., 2011). Recently, SA-mediated proline synthesis has been elucidated in relation to SA-dependent NPR1-mediated redox control with an antagonistic depression of ABA-signaling (La *et al*., 2019). Furthermore, Ca^2+^-dependent protein kinases (CPKs) are now known to play a central role in innate immune as a stress signaling by collaborating with hormonal signaling (Boudsocq and Sheen, 2013; Seyfferth and Tsuda, 2014). However, the ambivalent roles of ROS and proline in promoting stress tolerance and developing hypersensitive toxicity in connection with hormonal signaling pathway remain poorly understood.

Accordingly, the aims of the present study were to investigate the following hypotheses: 1) that exogenous Glu-application would enhance proline synthesis and subsequently modify the interplay between ROS and proline metabolism in association with hormonal regulation under drought stress, and 2) that stress response and tolerance mechanisms are differently regulated by the modified hormonal state and their signaling. To test these hypotheses, the drought-responsive hormonal status, ROS production, proline metabolism, and redox state were compared to the exogenous Glu-mediated changes with intention of characterizing hormonal regulation.

## Materials and Methods

### Plant growth and treatment

*Brassica napus* (cv.Pollen) seeds were germinated in the bed soil in a tray. Upon reaching the four-leaf stage, seedlings were transplanted in 2 litter pots that contained a 70:30 (w:w) mixture of soil and perlite, and grown with 100 ml nutrients solution (Lee *et al*., 2015). At the 6-7 leaves stage, plants were selected by morphological similarity and divided into two groups for the drought treatment. One group was normally irrigated with 200 ml for well-watered plants or with 20 ml for drought-stressed plants. After 5 days of drought treatment, both the well-watered and drought-treated group were divided into two sub-groups that were applied without or with 20 mM glutamate for 10 days. Glutamate application was done on the basis of preliminary test referring to the previous study (Kan *et al*., 2017). Thus, the experiment consisted of 4 treatments: well-watered (Control), Glu-application under well-watered (Glu), drought alone (Drought), and Glu-application under drought condition (Drought + Glu). The plants were grown in a greenhouse with day/night mean temperature of 27/20 °C and relative humidity 2 of 65/85%. Natural light was supplemented by metal halide lamps that generated 200 μmol photons m^-2^ s^-1^ at the canopy height for 16 h per day. The sampling was conducted at 5 days of drought (d 5) and at 10 days after glutamate application (d 15), respectively.

### Osmotic potential and measurement of photosynthetic pigment content

For the measurement of osmotic potential, fresh leaves were frozen in liquid nitrogen and then allowed them to thaw, followed by centrifugation at 13,000 ×*g* for 15 min. The collected sap was used for measuring osmolality by using a vapor pressure osmometer (Wescor 5100; Wescor Inc., Logan, UT). For total chlorophyll and carotenoid content, fresh leaves (100 mg) were immersed in 10 ml of 99% dimethyl sulfoxide. After 48 h, the absorbance of the supernatants was read at 480 nm and 510 nm for carotenoid, and 645 nm and 663 nm for total chlorophyll by using a microplate reader (Synergy H1 Hybrid Reader; Biotek, Korea).

### Determination of ROS production and antioxidative enzymes activity

For the visualization of H_2_O_2_ and O_2_^·–^, leaf discs were stained with 3,3’-diaminobenzidine (DAB) and nitroblue tetrazolium (NBT), respectively, as described previously (Lee *et al*., 2009a; Islam *et al*., 2017). The activity of superoxide dismutase (SOD) and catalase (CAT) were determined using the method of Lee *et al*. (2013). One unit of SOD enzyme activity was defined as the amount of enzyme required to inhibit 50% of the NBT photoreduction observed in negative control reactions. One unit of CAT enzyme activity was defined as the amount of enzyme required to degrade 1 mM H_2_O_2_ min^-1^.

### Measurement of cytosolic Ca^2+^ concentration

Cytosolic Ca^2+^ levels were estimated using aequorin luminometry detection (Tanaka *et al*., 2010) with some modifications. Briefly, 200 mg fresh leaves were extracted in a buffer solution containing 1 mM KCl, 1 mM CaCl_2_, and 10 mM MgCl_2_, adjusted pH to 5.7 using Tris-base, and centrifuged at 12,000 *×g* for 10 min. One hundred micro litter of supernatant was incubated with 1 μl of 0.1 mM coelenterazine-h in a 96-well plate for 30 min to facilitate binding between coelenterazine-h (Sigma) and aequorin. After incubation, an equal volume of 2 M CaCl_2_, which was dissolved in 30% ethanol (v/v), was added to discharge the remaining aequorin. Calcium concentration was determined by luminescence, according to Knight *et al*. (1996).

### Determination of proline and Δ^1^-pyrroline-5-carboxylate content

For the determination of proline and pyrroline-5-carboxylate (P5C) content, fresh leaf (200 mg) was homogenized in 3% sulfosalicylic acid and centrifuged at 13,000 *×g* for 10 min. The resulting supernatants were mixed with ninhydrin solution containing acetic acid and 6 M H_3_PO_4_ (v/v, 3:2) and boiled at 100 °C for 1 h. Then, toluene was added to the mixture, which was incubated for 30 min. The absorbance was determined at 520 nm and quantified as described previously (Lee *et al*., 2009b). P5C content was determined according to method described by Mezl and Knox (1976). The supernatants were mixed with 10 mM of 2-aminobenzaldehyde dissolved in 40% ethanol. Then, the mixture was incubated at 37 °C for 2 h to develop the yellow color. The absorbance was measured at 440 nm and calculated by using an extinction coefficient 2.58 mM^-1^ cm^-1^.

### Collection of phloem exudate and xylem sap

Phloem exudates were collected in ethylenediaminetetraacetic acid (EDTA) using the facilitated diffusion method, as described previously (Lee *et al*., 2009b). The fourth fully extended leaf was cut and immediately rinsed in 20 mM EDTA solution (pH 7.0) for 5 min. The leaf was then transferred to a new tube containing 5 mM EDTA solution and kept for 6 h in a growth chamber with 95% relative humidity under dark conditions. Xylem sap was collected by a vacuum-suction technique (Kotov and Kotova, 2015). Both the phloem exudates and xylem sap were stored at −20°C for further analysis.

### Measurement of glutathione and pyridine nucleotides

For the extraction of glutathione, 200 mg fresh leaves were homogenized in 5% 5-sulfosalicylic acid and centrifuged at 12,000 *×g* for 10 min. The glutathione content of the resulting supernatants was then determined by microplate assay using the GSH/GSSG Kit GT40 (Oxford Biomedical Research, Inc.). The contents of oxidized and reduced pyridine nucleotides were measured as described previously (La *et al*., 2019).

### Phytohormone analysis

Quantitative analysis of phytohormones in leaf tissue was performed by a high-performance liquid chromatography-electrospray ionization tandem mass spectrometry (HPLC–ESI–MS/MS) (La *et al*., 2019). Brief, fifty milligrams of fresh leaves in a 2-ml tube was frozen in liquid nitrogen and ground using a Tissuelyser II (Qiagen). The ground sample was extracted with 500 μl of extraction solvent, 2-propanol/H_2_O/concentrated HCl (2:1:0.002, v/v/v). Dichloromethane (1 ml) was added to the supernatant, and this was then centrifuged at 13,000 *×g* for 5 min at 4 °C. The lower phase, which was poured into a clean screw-cap glass vial, was dried under nitrogen and dissolved in pure methanol. The completely dissolved extract, ensured by vortexing and sonicating, was transferred to a reduced volume liquid chromatography vial. Hormones were analyzed by a reverse phase C18 Gemini high-performance liquid chromatography (HPLC) column for HPLC–ESI–MS/MS analysis. The chromatographic separation of hormones and its internal standard from the plant extracts was performed on an Agilent 1100 HPLC (Agilent Technologies), Waters C18 column (15,092.1 mm, 5 l m), and API3000 MSMRM (Applied Biosystems), using a binary solvent system comprising 0.1% formic acid in water (Solvent A) and 0.1% formic acid in methanol (Solvent B) at a flow rate of 0.5 ml/min.

### RNA extraction and quantitative real-time PCR analysis

Total RNA was isolated from 200 mg fresh leaf using an RNAiso Plus (Takara, DALIAN), and cDNA was synthesized using the GoScript Reverse Transcription System (Promega). Gene expression was quantified using a light cycle real-time PCR detection system (Bio-Rad, Hercules, CA, USA) with SYBR Premix Ex Taq (Takara, DALIAN). The PCR reactions were performed using the following conditions: 95 °C for 5 min; and then followed by 45 cycles of 95 °C for 30 s, 55–60 °C for 30 s, and 72 °C for 30 s; and a final extension of 72 °C for 5 min. The qRT-PCR was performed using genespecific primers (Supplementary Table S1). The qPCR reactions were performed in triplicate for each of three independent samples, and the relative expression levels of the target genes were calculated from threshold values (Ct), using the 2^-ΔΔCT^ method (Livak and Schmittgen, 2001) and the actin gene as an internal control.

### Statistical analysis

The present study used a completely randomized design with three replicates for each treatment and sampling date. Analysis of variance (ANOVA) was applied to all data, and Duncan’s multiple range test was used to compare the means of separate replicates. All statistical tests were performed using SAS 9.1 (SAS Institute, Inc., 2002-2003), and differences at *P* < 0.05 were considered significant. The heatmap, correlation coefficient, and pathway impact analyses were performed using MetaboAnalyst 3.0 (http://www.metaboanalyst.ca).

## Results

### Physiological symptoms, osmotic potential, and pigments

Drought stress induced severe leaf wilting and reduction in leaf osmotic potential. However, drought-induced negative effects were diminished in the glutamate (Glu)-treated plants (Fig. 1A, B). Chlorophyll and carotenoid contents were less or significantly reduced, respectively, by drought stress; however, both were greatly enhanced by Glu treatment. Under the well-watered conditions, exogenous Glu treatment significantly enhanced the content of these two pigments (Fig. 1C, D).

**Fig 1.**
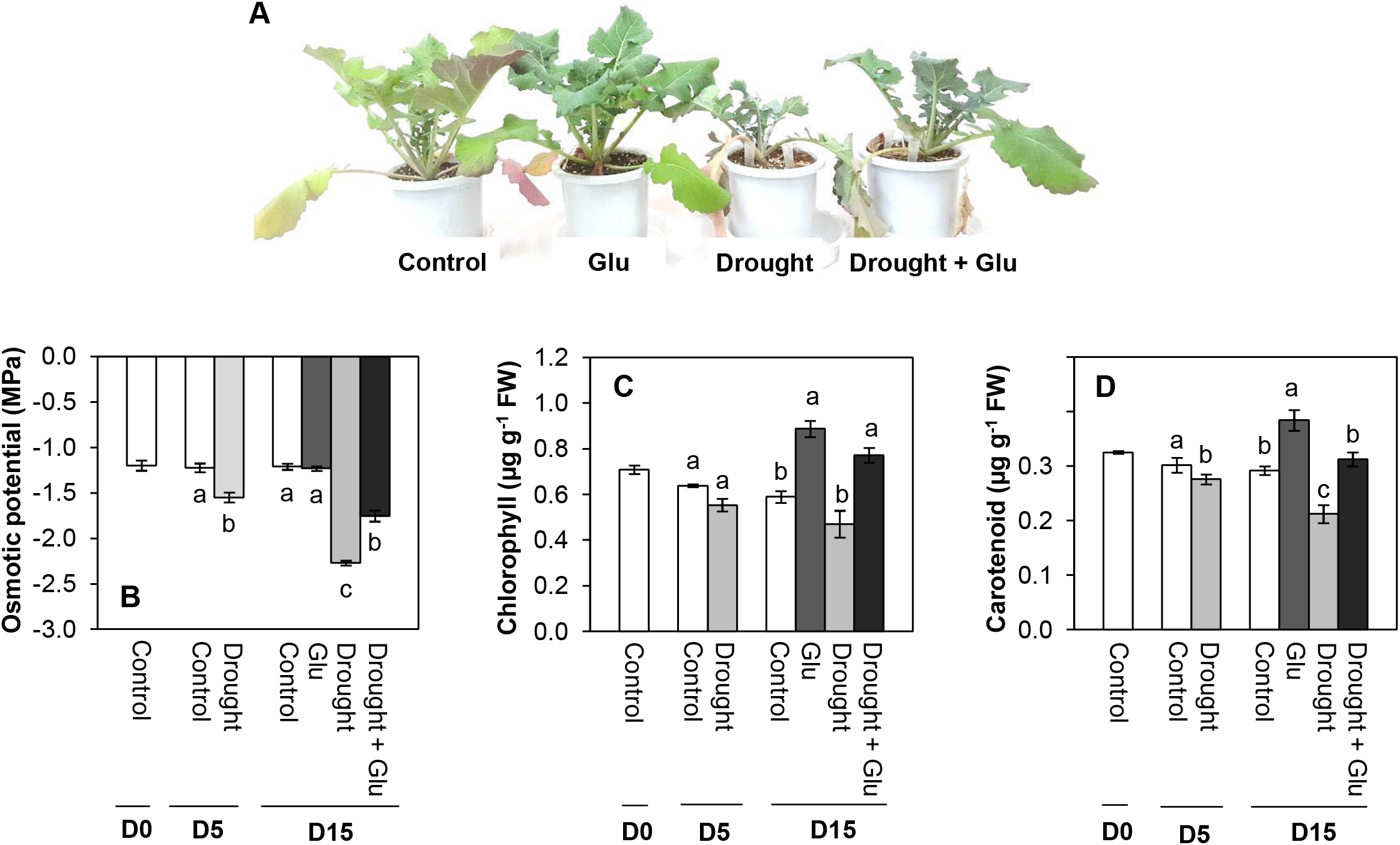
Changes in plant morphology, osmotic potential, and chlorophyll and carotenoid content in the leaves of control or glutamate (Glu)-treated *Brassica napus* under well-watered or drought-stressed conditions. (A) Plant morphology, (B) osmotic potential, (C) chlorophyll content, and (D) carotenoid content. Values are represented as mean ± SE (n = 3). Different letters on columns indicate significant difference at *P* < 0.05 according to the Duncan’s multiple range test.

### Phytohormone content and related gene expression

Endogenous level of abscisic acid (ABA) was remarkably increased in the treatment of drought alone (6.4-fold higher than that in the control), whereas it was significantly alleviated in the Drought + Glu treatment (3.6-fold higher than that in the control) at day 15. In contrast, drought-induced salicylic acid (SA) accumulation was further elevated in the Drought + Glu treatment (20% higher than that in drought alone). No significant difference was observed in the Glu treatment under well-watered conditions (Fig. 2A, B). In drought-stressed plants at day 15, compared to the control, content of IAA (indole-3-acetic acid) and CK (cytokinin) were largely increased by 2.1- and 1.3-fold, respectively, regardless of Glu treatment. IAA content was significantly increased in Glu-treated plants under the well-watered condition, whereas CK content was largely reduced (Fig. 2C, D). These results coincided with the expression pattern of hormone synthesis or signaling regulatory genes (Fig. 3).

**Fig 2.**
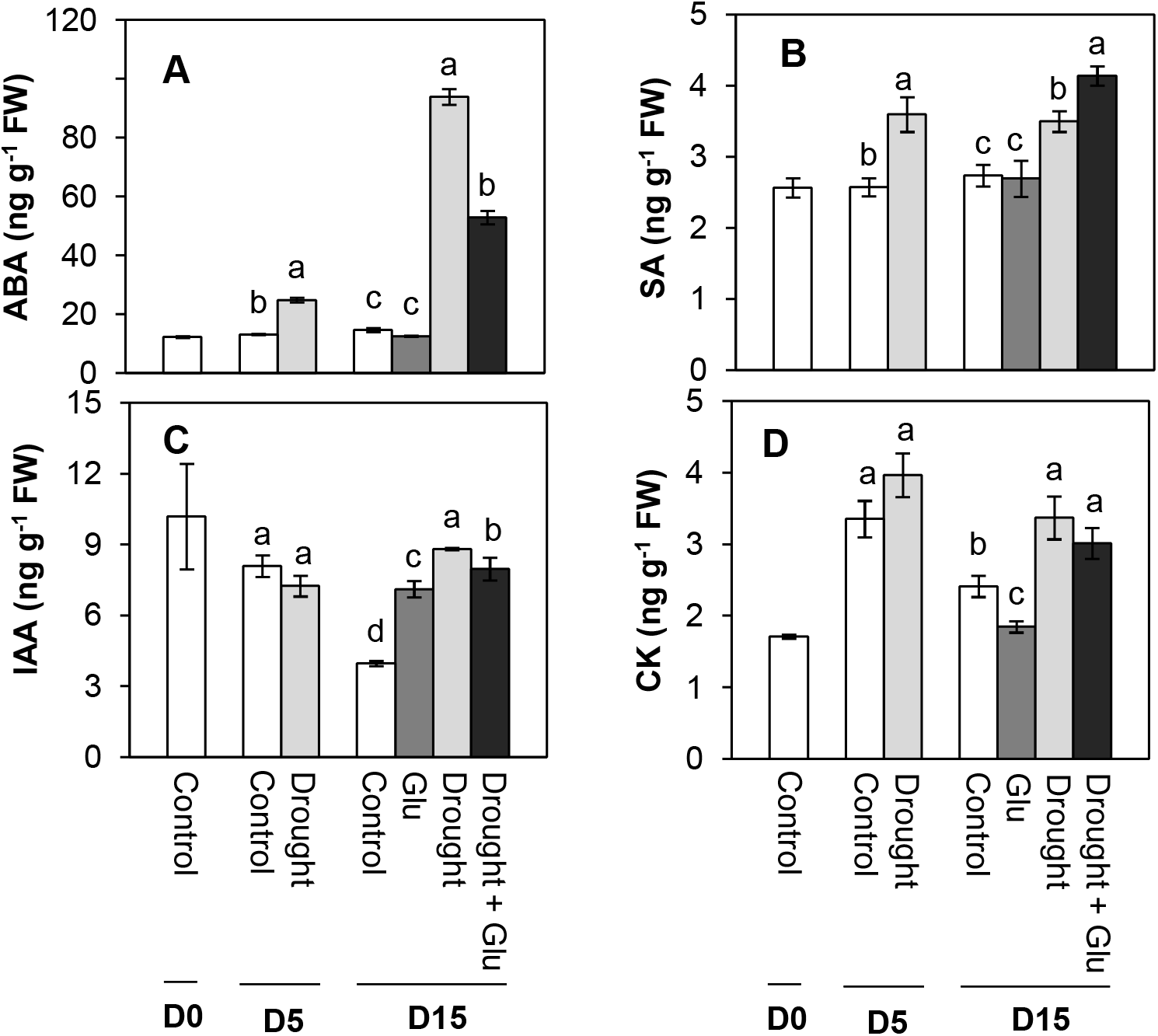
Phytohormone content in the leaves of control or glutamate (Glu)-treated *Brassica napus* under well-watered or drought stress conditions. (A) Abscisic acid (ABA), (B) salicylic acid (SA), (C) indole-3-acetic acid (IAA), and (D) cytokinin (CK) content. Values are represented as mean ± SE (n = 3). Different letters on columns indicate significant difference at *P* < 0.05 according to the Duncan’s multiple range test.

Drought stress remarkably upregulated the expression of the ABA signaling-related genes, myb-like transcription factor (*MYB2.1*) and NAC domain-containing protein 55 (*NAC55)*. However, enhanced expression of these two genes was largely depressed by the Drought + Glu treatment (Fig. 3A, B). In addition, expression of the SA synthesis-related genes, WRKY transcription factor 28 (*WRKY28*) and isochorismate synthase 1 (*ICS1*), were significantly upregulated by drought. A much higher expression of these genes was observed in the Drought + Glu treatment (Fig. 3C, D). Expression of the SA signaling related genes, nonexpressor of pathogenesis-related (PR) gene (*NPR1*) and *PR-1*, were significantly depressed upon drought stress at day 5 and, then, significantly upregulated at day 15. The Drought + Glu treatment further upregulated the expression of *NPR1* and *PR1* (Fig. 3E, F). No significant difference in these genes was observed in the Glu treatment under the well-watered conditions, expect for *NPR1* and *PR1* (Fig. 3A-F).

**Fig 3.**
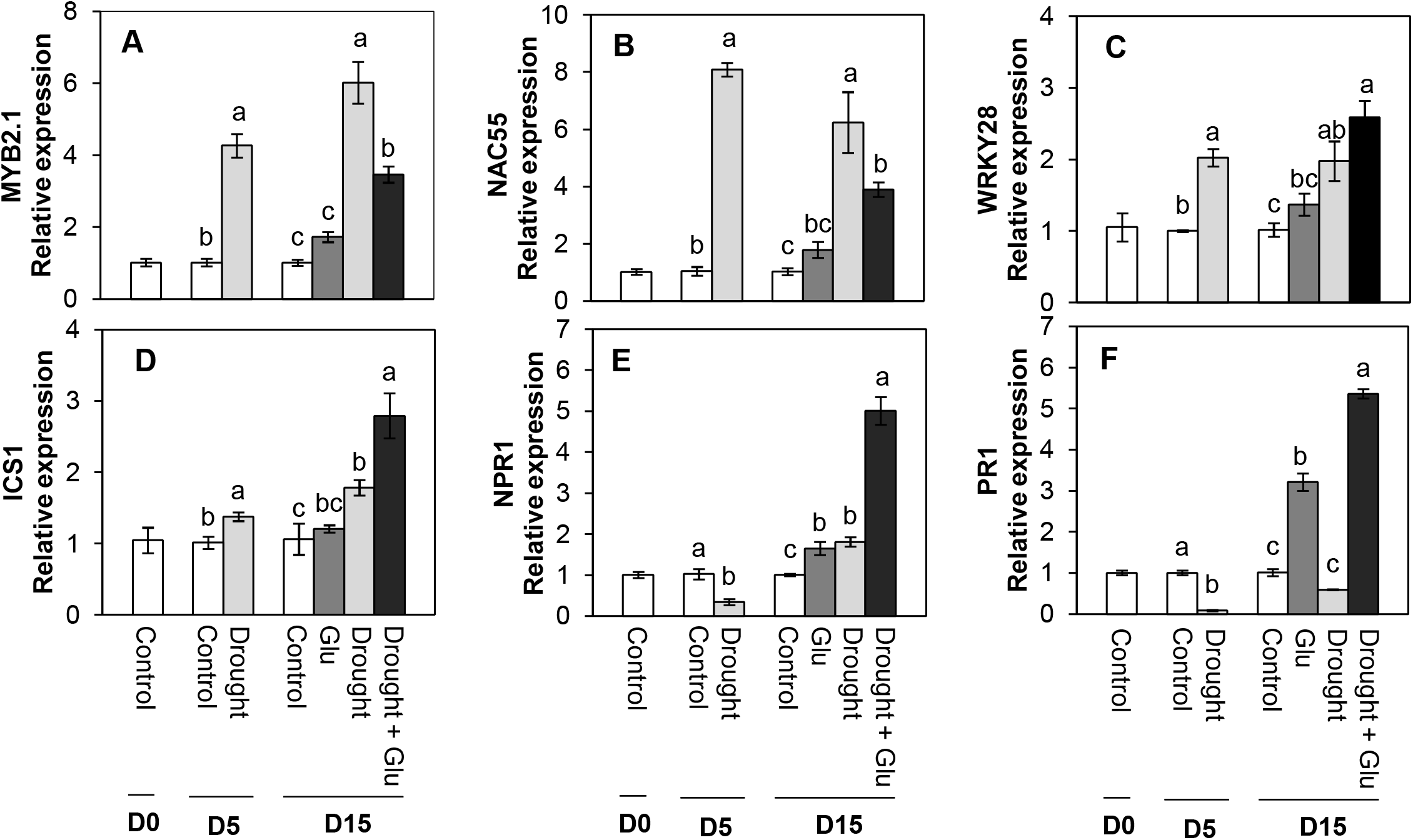
The relative expression of genes related to phytohormone synthesis or signaling in the leaves of control or glutamate (Glu)-treated *Brassica napus* under well-watered or drought-stressed conditions. ABA-responsive genes (A) myb-like transcription factor (*MYB2.1*) and (B) NAC domain-containing protein 55 (*NAC55*); SA synthesis-related genes (C) WRKY transcription factor 28 (*WRKY28*) and (D) isochorismate synthase 1 (*ICS1);* SA-responsive genes (E) nonexpressor of pathogenesis-related (PR) gene (*NPR1*) and (F) *PR-1*. qRT-PCR was performed in duplicate for each of the three independent biological samples. Values are represented as mean ± SE (n = 3). Different letters on columns indicate significant difference at *P* < 0.05 according to the Duncan’s multiple range test.

### Glutamate receptor, ROS, Ca^2+^ signaling, and antioxidant activity

The expression of glutamate receptor, GLR1.3, was remarkably upregulated by drought stress. It was enhanced considerably by Glu treatment (Fig. 4A). A significant accumulation of ROS (O_2_ and H_2_O_2_) production was observed with *in situ* localization of O_2_ and H_2_O_2_ under drought treatment, indicated by dark spots (Fig. 4B, C). Cytosolic Ca^2+^ content significantly increased with drought treatment, with 56% in the drought alone treatment and 85% in the Drought + Glu treatment compared to that in the control (Fig. 4D). Expression of calcium signaling-related gene, calcium-dependent protein kinase 5 (*CPK5*) was significantly induced by drought and/or Glu treatments throughout the experimental period. The greatest level was observed in the Drought + Glu treatment (Fig. 4E). The expression of *NADPH oxidase* was only enhanced significantly with drought alone treatment (Fig. 4F). Superoxide dismutase (SOD) activity was largely increased under drought conditions, regardless of Glu treatment (Supplementary Fig. S1A). The drought-induced increase in catalase (CAT) activity and its gene expression was further activated by Glu treatment (Supplementary Fig. S1B, C).

**Fig 4.**
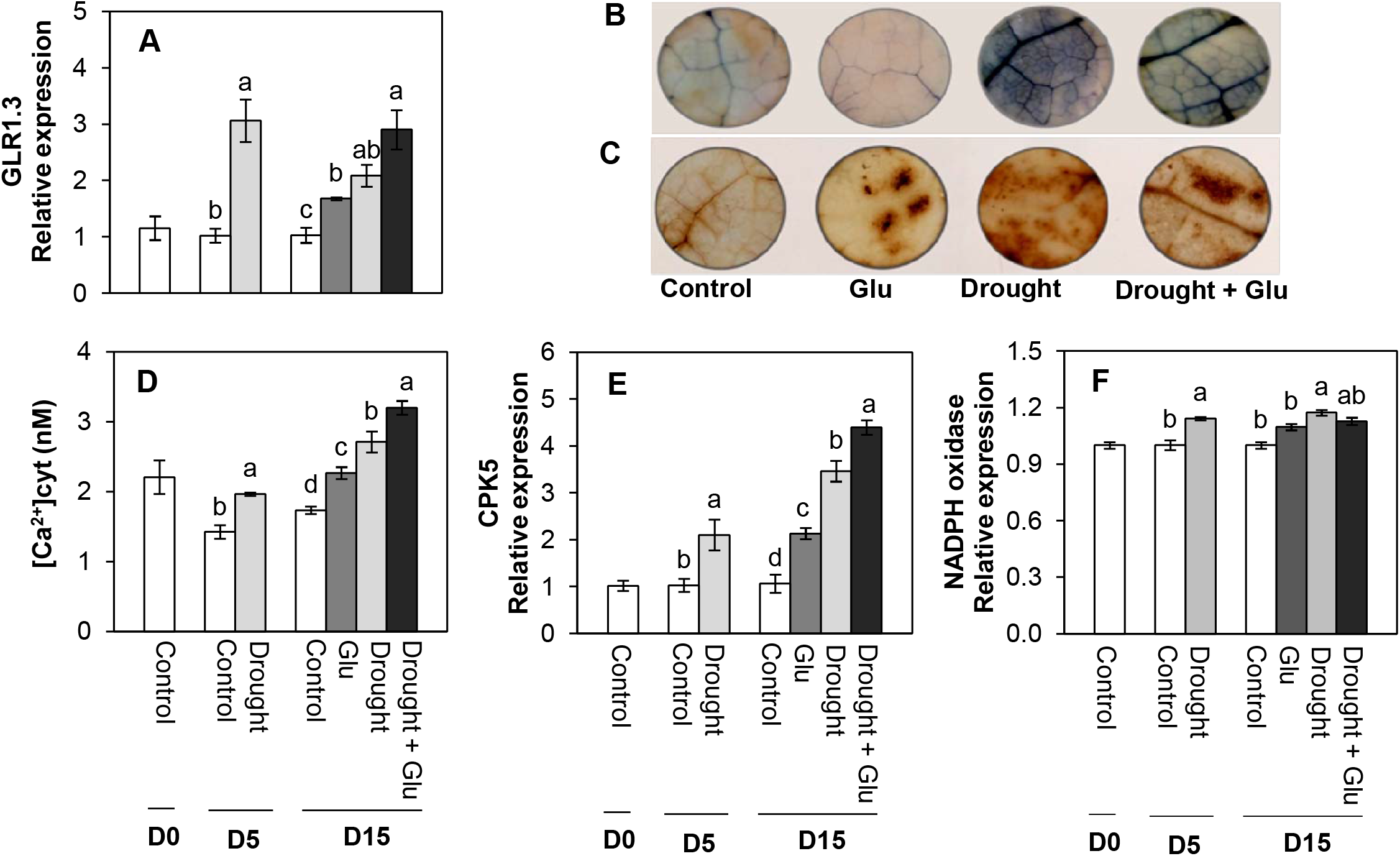
Changes in glutamate receptor, ROS, Ca^2+^ content and its signaling, and NADPH oxidases in the leaves of control or glutamate (Glu)-treated *Brassica napus* under well-watered or drought-stressed conditions. (A) Glutamate receptor (*GLR1.3)*, visualization of (B) O_2_^-^ and (C) H_2_O_2_, (D) Ca^2+^ content, (E) calcium-dependent protein kinase 5 (*CPK5*), and (F) *NADPH oxidase*. qRT-PCR was performed in duplicate for each of the three independent biological samples. Values are represented as mean ± SE (n = 3). Different letters on columns indicate significant difference at *P* <0.05 according to the Duncan’s multiple range test.

### Proline metabolism and transport

Pyrroline-5-carboxylate (P5C) content was significantly increased by 2.4-fold under drought with or without Glu treatment at day 15, compared with that in the control (Fig. 5C). Drought stress significantly induced proline accumulation throughout the experimental period, with a much greater increase in the Drought + Glu treatment (2.7 fold-higher than that in the drought alone treatment; Fig. 5E). Expression of proline synthesis-related genes, P5C synthase 1 (*P5CS1*), *P5CS2*, and P5C reductase (*P5CR*), were remarkably upregulated by drought and/or Glu treatment. Expression of these genes was much higher in the Drought + Glu treatment (Fig. 5A, B, D). The proline degradation-related genes, proline dehydrogenase (*PDH*) and pyrroline-5-carboxylate dehydrogenase (*P5CDH)*, were differently expressed during the experimental period. The expression of *PDH* was largely depressed by drought and/or Glu treatments, whereas expression of *P5CDH* was significantly enhanced by the drought treatment (Fig. 5F, G). Proline content in the phloem and xylem was greatly increased by drought and/or Glu treatments. The highest proline content was observed in the Drought + Glu treatment (Supplementary Fig. S2A, B).

**Fig 5.**
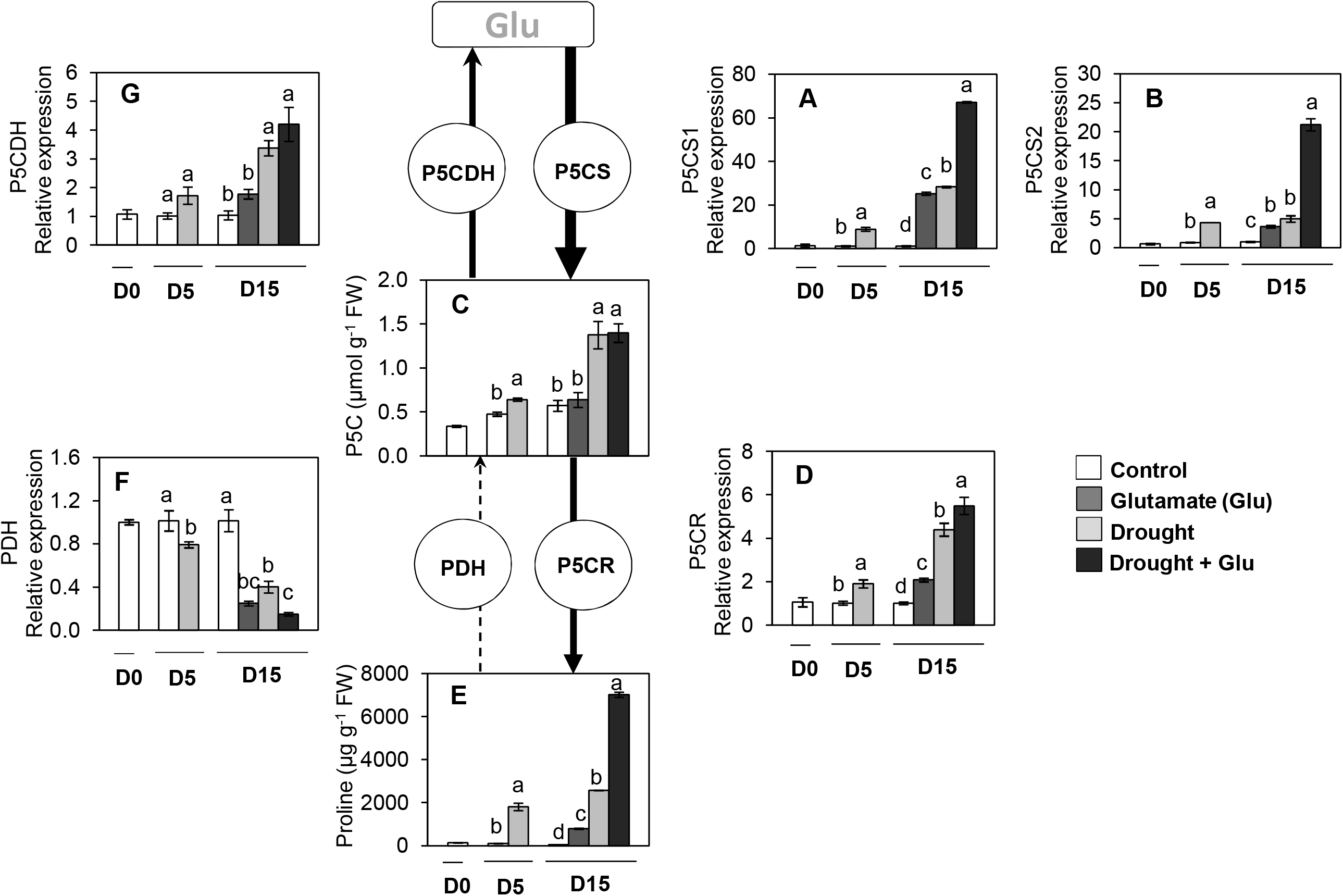
Changes in proline metabolism in the leaves of control or glutamate (Glu)-treated *Brassica napus* under well-watered or drought-stressed conditions. (A) Pyrroline-5-carboxylate (P5C) synthase 1 (*P5CS1*), (B) *P5CS2*, (C) P5CS content, (D) pyrroline-5-carboxylate reductase (*P5CR*), (E) proline content, (F) proline dehydrogenase (*PDH)*, and (G) pyrroline-5-carboxylate dehydrogenase (*P5CDH)*. qRT-PCR was performed in duplicate for each of the three independent biological samples. Values are represented as mean ± SE (n = 3). Different letters on columns indicate significant difference at *P* < 0.05 according to the Duncan’s multiple range test.

### Redox status and redox signaling component

NAD(P)H content was significantly increased by drought stress compared with that of the control, whereas it was much more enhanced by Glu treatment. Drought-induced NAD(P)^+^ accumulation was significantly alleviated by Glu treatment. The ratio of NAD(P)H to NAD(P)^+^ was largely decreased by the drought treatment. However, its reduction was largely mitigated in the Drought + Glu treatment. Reduced glutathione (GSH) content was greatly decreased by 86.3% under the drought alone treatment compared with that under the control, whereas it recovered to 72.7% of that in the control in the Drought + Glu treatment. Oxidized glutathione (GSSG) content was similar between treatments. Drought-induced reduction of the ratio of GSH to GSSG was largely recovered with Glu treatment (Table 1). Drought and/or Glu treatments significantly enhanced the expression of the oxidoreductase-encoding genes, CC-type glutaredoxin 9 (*GRXC9*) and thioredoxin-h5 (*TRXh5*), and this increase was much higher in the Drought + Glu treatment (Fig. 6A, B). The expression of TGA-box transcription factor (*TGA2*) was upregulated only in the Drought + Glu treatment (Fig. 6C).

**Fig 6.**
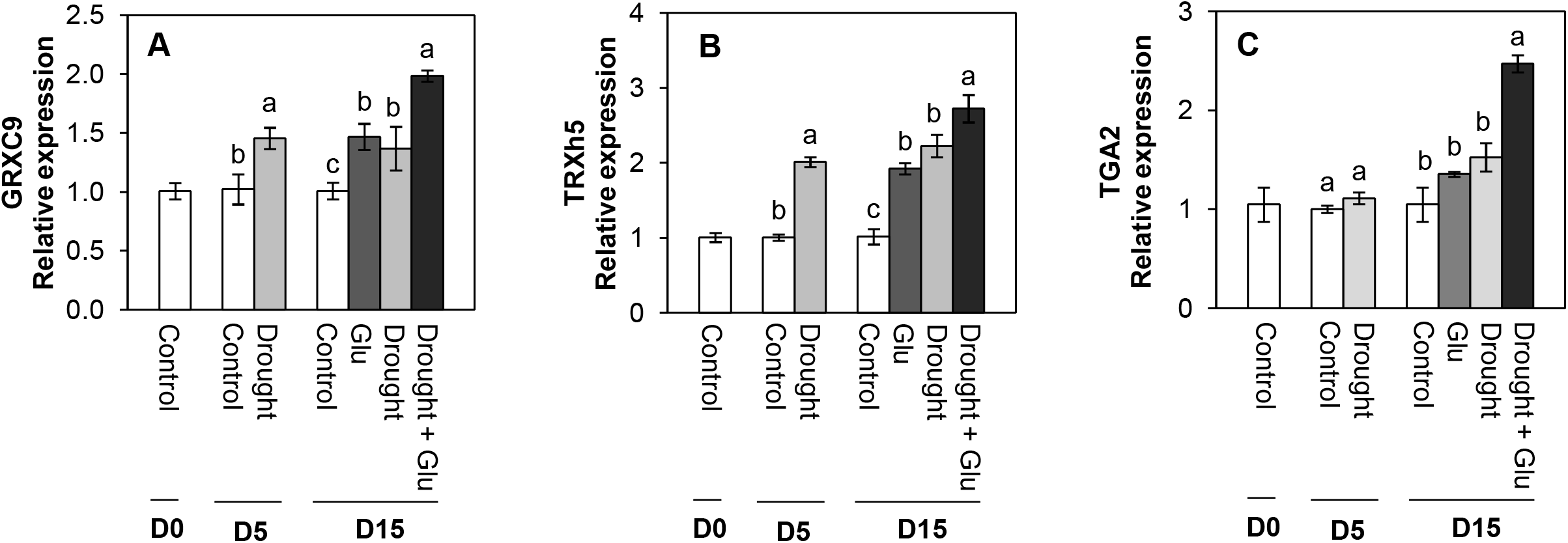
Relative expression of genes related to redox signaling in the leaves of control or glutamate (Glu)-treated *Brassica napus* under well-watered or drought-stressed conditions. (A) CC-type glutaredoxin 9 (*GRXC9*), (B) thioredoxin-h5 (*TRXh5*), and (C) TGA-box transcription factor (*TGA2*). qRT-PCR was performed in duplicate for each of the three independent biological samples. Values are represented as mean ± SE (n = 3). Different letters on columns indicate significant difference at *P* < 0.05 according to the Duncan’s multiple range test.

**Table 1.** Changes in redox status in the leaves of control or glutamate (Glu)-treated *Brassica napus* under well-watered or drought-stressed conditions

| Treatments | Days after treatment |  |  |  |  |  |  |  |  |
| --- | --- | --- | --- | --- | --- | --- | --- | --- | --- |
|  | 0 |  |  | 5 |  |  | 15 |  |  |
| Reduced | NADPH | NADH | GSH | NADPH | NADH | GSH | NADPH | NADH | GSH |
| Control | 2.65 ± 0.83 | 5.39 ± 0.48 | 61.51 ± 4.81 | 2.62 ± 0.17<sup>b</sup> | 5.58 ± 0.44<sup>a</sup> | 58.58 ± 2.01<sup>a</sup> | 2.32 ± 0.25<sup>c</sup> | 4.44 ± 0.33<sup>c</sup> | 53.20 ± 3.26<sup>a</sup> |
| Glu | - | - | - | - | - | - | 5.28 ± 0.39<sup>a</sup> | 5.07 ± 0.40<sup>bc</sup> | 60.35 ± 5.00<sup>a</sup> |
| Drought | - | - | - | 3.48 ± 0.25<sup>a</sup> | 5.79 ± 0.12<sup>a</sup> | 15.77 ± 0.21<sup>b</sup> | 3.78 ± 0.10<sup>b</sup> | 5.37 ± 0.03<sup>b</sup> | 7.27 ± 0.12<sup>c</sup> |
| Drought + Glu | - | - | - | - | - | - | 4.84 ± 0.04<sup>b</sup> | 6.56 ± 0.03<sup>a</sup> | 38.67 ± 1.87<sup>b</sup> |
| Oxidized | NADP<sup>+</sup> | NAD<sup>+</sup> | GSSG | NADP<sup>+</sup> | NAD<sup>+</sup> | GSSG | NADP<sup>+</sup> | NAD<sup>+</sup> | GSSG |
| Control | 8.86 ± 0.06 | 6.81 ± 0.06 | 2.30 ± 0.05 | 9.62 ± 0.59<sup>b</sup> | 6.82 ± .30<sup>b</sup> | 2.62 ± 0.05<sup>b</sup> | 7.23 ± 1.05<sup>d</sup> | 6.01 ± 0.22<sup>c</sup> | 2.45 ± 0.09<sup>a</sup> |
| Glu | - | - | - | - | - | - | 14.36 ± 0.61<sup>c</sup> | 8.53 ± 0.25<sup>b</sup> | 3.10 ± 0.25<sup>a</sup> |
| Drought | - | - | - | 17.80 ± 1.00<sup>a</sup> | 9.10 ± 0.17<sup>a</sup> | 3.23 ± 0.13<sup>a</sup> | 24.57 ± 0.30<sup>a</sup> | 9.89 ± 0.07<sup>a</sup> | 2.69 ± 0.22<sup>a</sup> |
| Drought + Glu | - | - | - | - | - | - | 21.36 ± 0.52<sup>b</sup> | 9.51 ± 0.17<sup>a</sup> | 2.47 ± 0.22<sup>a</sup> |
| Ratios | NADPH/<br>NADP<sup>+</sup> | NADH/<br>NAD<sup>+</sup> | GSH/<br>GSSG | NADPH/<br>NADP<sup>+</sup> | NADH/<br>NAD<sup>+</sup> | GSH/<br>GSSG | NADPH/<br>NADP<sup>+</sup> | NADH/<br>NAD<sup>+</sup> | GSH/<br>GSSG |
| Control | 0.41 ± 0.02 | 0.79 ± 0.07 | 26.65 ± 1.98 | 0.26 ± 0.00<sup>a</sup> | 0.81 ± 0.03<sup>a</sup> | 25.94 ± 1.02<sup>a</sup> | 0.27 ± 0.02<sup>b</sup> | 0.74 ± 0.06<sup>a</sup> | 21.70 ± 0.67<sup>a</sup> |
| Glu | - | - | - | - | - | - | 0.37 ± 0.02<sup>a</sup> | 0.60 ± 0.04<sup>ab</sup> | 19.58 ± 1.20<sup>ab</sup> |
| Drought | - | - | - | 0.21 ± 0.01<sup>a</sup> | 0.64 ± 0.02<sup>b</sup> | 4.92 ± 0.25<sup>b</sup> | 0.15 ± 0.03<sup>c</sup> | 0.54 ± 0.00<sup>b</sup> | 2.75 ± 0.21<sup>c</sup> |
| Drought + Glu | - | - | - | - | - | - | 0.21 ± 0.00<sup>bc</sup> | 0.69 ± 0.01<sup>ab</sup> | 15.96 ± 1.36<sup>b</sup> |

Reduced form of nicotinamide adenine dinucleotide (phosphate), NAD(P)H; oxidized form of nicotinamide adenine dinucleotide (phosphate), NAD(P)<sup>+</sup>; reduced form of glutathione, GSH; oxidized form of glutathione, GSSG. NAD(P)H and NAD(P)<sup>+</sup> contents are shown as nmol g<sup>-1</sup> fresh weight. GSH and GSSG contents are shown as μmol g<sup>-1</sup> fresh weight. Values are mean ± SE for n = 3. Different lowercase letters in a column indicate significant differences at *P* < 0.05 according to the Duncan's multiple range test.

### Heatmap visualization and Pearson correlation analysis for the metabolites or gene expression

To further clarify the metabolites or gene expression levels affected by the drought-stress and/or Glu treatments, the results of hormones, ROS, upstream ROS signal, glutamate receptor, proline metabolism, redox status, and their signaling were visualized by heatmap and Pearson correlation coefficients (Fig. 7). The drought effect was notably higher on the increase of ABA and its signaling gene *MYB2.1*, H_2_O_2_, NADPH oxidase as well as on the reduction of reducing potential [NAD(P)H/NAD(P)^+^ and GSH/GSSG]. These drought effects were alleviated with the Drought + Glu treatment, resulting in an increase in SA and its synthesis or signaling gene (*NPR1* or *WRKY28*, respectively), CPK5, reducing potential, and proline synthesis (Fig. 7A). The correlations of proline revealed a positive relation with the expression of the SA-signaling regulatory genes *NPR1* and *CPK5*. In addition, SA was closely correlated with the reducing power (Fig. 7B).

**Fig 7.**
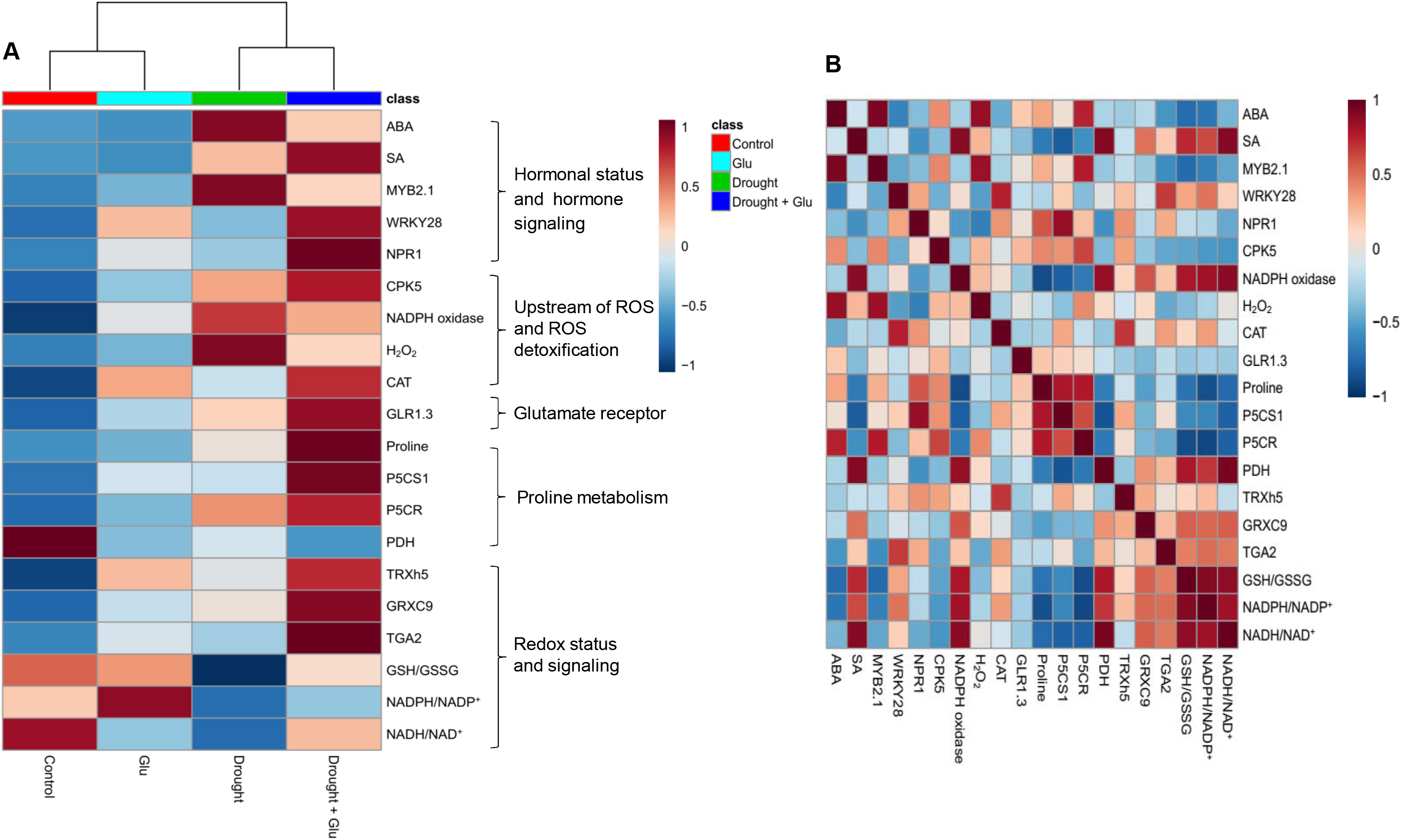
Heatmap analysis of the treatment effect and correlations among the variables measured at day 15 (after 15 d of drought, including 10 d of glutamate application). (A) Heatmap comparing the changes in the identified metabolites or gene expression levels in the leaves of control or glutamate (Glu)-treated plants under well-watered or drought-stressed conditions. The normalization procedure consisted of mean row-centering with color scales. (B) Heatmap showing the correlations among the identified metabolites or gene expression levels. Correlations coefficients were calculated based on Pearson’s correlation. Red indicates a positive effect, whereas blue indicates a negative effect. Color intensity is proportional to the correlation coefficients.

## Discussion

The accumulation of proline, which is considered as a representative compatible solute, is commonly observed in a wide range of abiotic and biotic stresses. This stress response is thought to function as a protective mechanism in stressed plants (Rejeb *et al*., 2014; Xia *et al*., 2015). However, proline metabolism is responsible for stress-induced ROS production and is, subsequently, involved in the hypersensitive response of plants (Liang *et al*., 2013). Therefore, determining the thresholds of regulatory mechanisms at which proline metabolism switches from hypersensitive responses to stress resistance (or *vice versa*) would provide valuable insight into the underlying mechanisms of plant stress responses. Accordingly, one of the aims of the present study was to test the hypothesis that exogenous Glu would accelerate proline synthesis, because proline is mainly synthesized from Glu under drought conditions (Rejeb *et al*., 2014) and because the early Glu-responsive genes encode membrane receptors, protein kinase/phosphatases, Ca^2+^ signaling, and transcription factors (Kan *et al*., 2017). The present study, thus, assessed preferentially the effect of Glu-responsive proline metabolism on drought symptom development.

In the present study, the 5-d drought treatment induced the accumulation of both ROS and proline, as is commonly observed in drought-stressed plants (Lee *et al*., 2009b; Rejeb *et al*., 2014; La *et al*., 2019), and another 10 d of drought (15 d in total) provoked severe drought symptoms, such as leaf wilting and reduced leaf osmotic potential (Fig. 1A, B). These drought-induced hypersensitive responses were accompanied with enhanced ROS accumulation (Fig. 4B, C) and reduced reducing potential (Table. 1). Severe drought symptom in drought alone reflected also the highest ABA accumulation and ABA-related genes expression (Fig. 2A and 3A, B). ABA has been reported to stimulate a signaling pathway that triggers ROS production, which in turn induces increases in cytosolic Ca^2+^ (Osakabe *et al*., 2014). Indeed, drought-induced ABA-mediated ROS accumulation was concomitant with increased levels of NADPH oxidase (Fig. 4F), accompanied by cytosolic Ca^2+^ (Fig. 4D) and CPK5 (Fig. 4E), which is consistent with the findings of previous studies (Boudsocq and Sheen, 2013; Rejeb *et al*., 2015; Stael *et al*., 2015). ROS (mainly H_2_O_2_) accumulation that is accompanied by redox changes might directly or indirectly involve in regulating the transcription of proline biosynthesis (Rejeb *et al*., 2015; La *et al*., 2019). In the present study, a significant accumulation of proline with enhanced expression of proline synthesis-related genes was observed in drought-stressed plants, regardless of Glu treatment (Fig. 5). Previous studies have also reported ABA-induced proline accumulation (Verslues *et al*., 2007). The simultaneous accumulation of ROS and ABA has been postulated as a key aspect of cross-tolerance (Verslues *et al*., 2007). Furthermore, the interplay between ABA, ROS and proline has been suggested to function as an integrative process in regulating water stress responses and signal transduction pathways (Verslues *et al*., 2007; Liang *et al*., 2013; Osakabe *et al*., 2014). However, in the present study, the drought-induced ABA-responsive enhancement of ROS and proline was a hypersensitive response that included the expression of severe symptoms, whereas the negative symptom induced by drought was significantly alleviated in the Drought + Glu treatment, despite the additional accumulation of ROS and proline (Figs 1, 4, and 5). It is, therefore, tempting to characterize the plant immune and stress-signaling networks that trigger appropriate and diverse downstream responses to drought stress. Of the many networks involved in responses to drought stress, the present study focused on Ca^2+^-dependent protein kinases (CPKs) because recent studies have highlighted the roles of CPK-signaling pathways in plant immune and stress responses (Boudsocq and Sheen, 2013; Stael *et al*., 2015; Prodhan *et al*., 2018). In the proposed model for interactions between ROS and Ca^2+^ signaling (Boudsocq and Shen, 2013; Stael *et al*., 2015), CPKs, upon activation by the Ca^2+^ flux, together with a mitogen-activated protein kinase (MAPK), trigger the expression of immunity-related genes (Stael *et al*., 2015). Meanwhile, several protein kinases, including CPKs, enhance the activity of Rbohs (i.e., NADPH oxidase), thereby promoting the generation of apoplastic ROS (Boudsocq and Shen, 2013; Dubiella *et al*., 2013). In the present study, the drought-stress treatment induced increases in glutamate receptor GLR1.3 (Fig. 4A), cytosolic Ca^2+^ (Fig. 4D), and *CPK5* expression (Fig. 4E), regardless of Glu treatment. Boudsocq and Sheen (2013) reported that the signal through ABA synthesis activates CPKs, which regulate ROS and proline accumulation, water transport (eg., aquaporin) as well as related genes expression. Indeed, in this study, the enhanced CPK5 expression in the treatment drought alone was concomitant with an accumulation of ROS (Fig. 4B, C) and proline (Fig. 5E), accompanied by the highest ABA level and ABA-signaling genes expression (Figs 2A and 3A, B). In rice, CPKs have been reported to enhance salt-stress tolerance by regulating ROS homeostasis through the induction of ROS scavenging genes (*APX2/APX3*) and the suppression of the NADPH oxidase gene, *Rboh1* (Asano *et al*., 2012). However, in the present study, drought-enhanced ABA-responsive CPK5 was not observed to either suppress NADPH oxidase or scavenge ROS (Fig. 4). Interestingly, the Drought + Glu treatment further up-regulated *CPK5* expression, thereby increasing both endogenous SA and the expression of SA synthesis- and signaling-related genes (*ICS1* and *NPR1*, respectively), with antagonistic depression of ABA level (Fig. 2A) and the expression of ABA-signaling genes (*MYB2.1* and *NAC55;* Fig. 3A, B). The increased SA and SA-related gene expression, which coincided with exogenous Glu-enhanced-CPK5, significantly reduced the accumulation of ROS (Fig. 4B, C) and increased the accumulation of proline (Fig. 5E), thereby alleviating the negative symptoms of drought stress (Fig. 1). It is worth noting that there was a remarkable difference in the drought symptoms of the Drought and Drought + Glu plants (Fig. 1A), even though plants in both treatments exhibited a significant accumulation of ROS and proline, as well as enhanced cytosolic Ca^2+^ and *CPK5* expression. These results demand further discussion of the hormonal regulatory pathways involved in the integrative process of stress tolerance, a discussion which should emphasize the most distinct differences in the hormonal balance and gene expression of the two treatment groups (Figs 2 and 3).

Several reviews have documented that ROS and proline that is accumulated in response to stress stimuli function as signaling molecules, with possible interactions with phytohormonal signaling in metabolic regulatory pathways (Szabados and Savoure’, 2010; Liang *et al*., 2013; Rejeb *et al*., 2014; Herrera-Vásquez *et al*., 2015). In the present study, the simultaneous and significant accumulation of ROS and proline, accompanied by elevated cytosolic Ca^2+^ and *CPK5* expression, was observed under drought stress, regardless of Glu treatment. However, the pattern of ROS and proline, as well as cytosolic Ca^2+^ and *CPK5* expression followed by ABA-dependent in the treatment Drought alone, while SA-dependent manner in the treatment Drought + Glu (Figs 2, 4, and 5A). Furthermore, drought-induced proline was much more increased in the treatment Drought + Glu, accompanied by further enhancements of proline synthesis-related genes (*P5CS* and *P5CR*) and depression of proline degradation-related gene (*PDH;* Fig. 5) expression. The accumulation of proline in response to exogenous Glu treatment, along with the additional activation of Ca^2+^ and CPK5, was induced in a SA-dependent manner (Figs 2B and 4D, E). The Ca^2+^-binding transcription factor CBP60g regulates the transcription of SA biosynthesis genes (e.g., *ICS1/SID2;* Zhang *et al*., 2010; Wang *et al*., 2011), thereby providing a venue for the Ca^2+^ signal to activate the WRKY28 transcription factor (Fig. 3C) in SA production. Indeed, the highest expression levels of *ICS1, NPR1*, and *PR1* in the Drought + Glu plants were consistent with the highest proline level and enhanced expression of proline synthesis-related genes (Figs 3D-F and 5), as well as with the downregulation of ABA (Fig. 2A). Similarly, Chen *et al*. (2011) reported that exogenous proline significantly induced intracellular Ca^2+^ accumulation and Ca^2+^-dependent ROS production, thereby enhancing SA synthesis. The results of several other studies have supported the interplay between SA and proline in regulating stress responses, e.g., proline-activated SA-induced protein kinase SIPK (Elizabeth and Zhang, 2000), involvement of SA in exogenous proline-induced salt resistance (Chen *et al*., 2011), and proline-mediated drought tolerance (La *et al*., 2019). Furthermore, elevated SA levels suppressed ROS production in the present study (Fig. 4B, C), potentially through a feedback loop for O_2_^·–^ (Straus *et al*., 2010) and the enhanced activation of CAT for scavenging H_2_O_2_ (Supplementary Fig. S1B). Indeed, SA-activated CAT (Herrera-Vásquez *et al*., 2015; La *et al*., 2019) and Ca^2+^-dependent CAT activation (Mou *et al*., 2003) have been reported previously. In addition, as far as we know, this study provides the first report of exogenous Glu-increased proline loading to both the xylem and phloem (Supplementary Fig. S2). Given that glutamate triggers long-distance, Ca^2+^-based plant defense signaling, it is reasonable to conclude that the Glu-mediated overproduction of proline could be responsible for SA production and the activation of SA-signaling and involve also in activation of Ca^2+^-mediated signaling, thereby functioning as a crucial regulatory pathway of stress tolerance. However, the mechanism by which proline- or SA-elicited ROS signals activate CPK5 remains unclear and requires further investigation.

Calcium-mediated signaling that occurs after the accumulation of SA has been reported to contribute to the regulation of defense-related gene expression. The interaction of Ca^2+^ is enhanced by the binding of Ca^2+^ to leucine zipper transcription factor TGA (Szymanski *et al*., 1996), which interacts with NPR1, a critical transcription cofactor in SA perception and the SA-mediated transcriptional regulation of PR1 through NPR1 (Seyfferth and Tsuda, 2014; Herrera-Vásquez *et al*., 2015), thereby providing a possible SA-mediated option to regulate stress tolerance reactions. In the present study, exogenous Glu-responsive, SA-mediated *NPR1* and *PR1* expression was consistent with the expression of TGA2 and CPK5, which was greatest in the Drought + Glu plants (Figs 3E-F, 4E, and 6C). Moreover, a synergistic and significant interaction between proline and SA for SA-transduction signaling (*NPR1* and *PR1*) was also observed in the Drought + Glu plants (Figs 3E-F and 5E). Increasing evidence demonstrates that *NPR1* is the first redox sensor to be described for SA-regulated genes and that *NPR1* is the master co-activator of *PR1* (Mou *et al*., 2003; Tada *et al*., 2008; Kneeshaw *et al*., 2014; La *et al*., 2019). Over-produced proline also activated the SA-signaling pathway but not the JA-signaling pathway (Chen *et al*., 2011).

Given that proline metabolism is directly control NAD(P)^+^/NAD(P)H redox balance (Sharma *et al*., 2011; Rejeb *et al*., 2014). These suggest that a significant recovery of reducing potential [GSH/GSSG and NAD(P)H/NAD(P)^+^ ratios] in the treatment Drought + Glu (Table 1) would be closely related with Glu-enhanced proline synthesis in a SA-mediated redox regulation. Given that proline metabolism is directly controlled by the NAD(P)^+^/NAD(P)H redox balance (Sharma *et al*., 2011; Rejeb *et al*., 2014), a significant recovery of reducing potential, i.e., GSH/GSSG and NAD(P)H/NAD(P)^+^ ratios, in response to the Drought + Glu treatment (Table 1) would be closely related to Glu-enhanced proline synthesis, as part of SA-mediated redox regulation. Indeed, in the Drought + Glu treatment, the oxidoreductase-encoding genes *TRXh5* and *GRXC9* were upregulated in a SA-mediated, NPR1-dependent manner (Figs 3E and 6). These genes are essential for redox control in SA-mediated transcriptional responses (Mou *et al*., 2003; Tada *et al*., 2008; Herrera-Vásquez *et al*., 2015). Therefore, the results of both the present study and previous reports (Mou *et al*., 2003; Seyfferth and Tsuda, 2014; La *et al*., 2019) provide evidence that SA-mediated, NPR1-dependent transcriptional responses, which may interact with proline metabolism, are integrative cellular redox regulation processes that promote PR1 induction.

The results of the heatmap and Pearson correlation analysis (Fig. 7) provide a basis for a working model of the signaling pathway that is activated by exogenous Glu (Fig. 8). In summary, the drought-induced negative stress responses were largely alleviated by exogenous Glu-induced, SA-mediated modulations that were characterized by 1) antagonistic depression of ABA-dependent metabolic and signaling pathways, 2) synergetic interaction of CPK5-mediated SA induction and proline synthesis, and 3) SA-mediated NPR1-dependent redox regulation.

**Fig 8.**
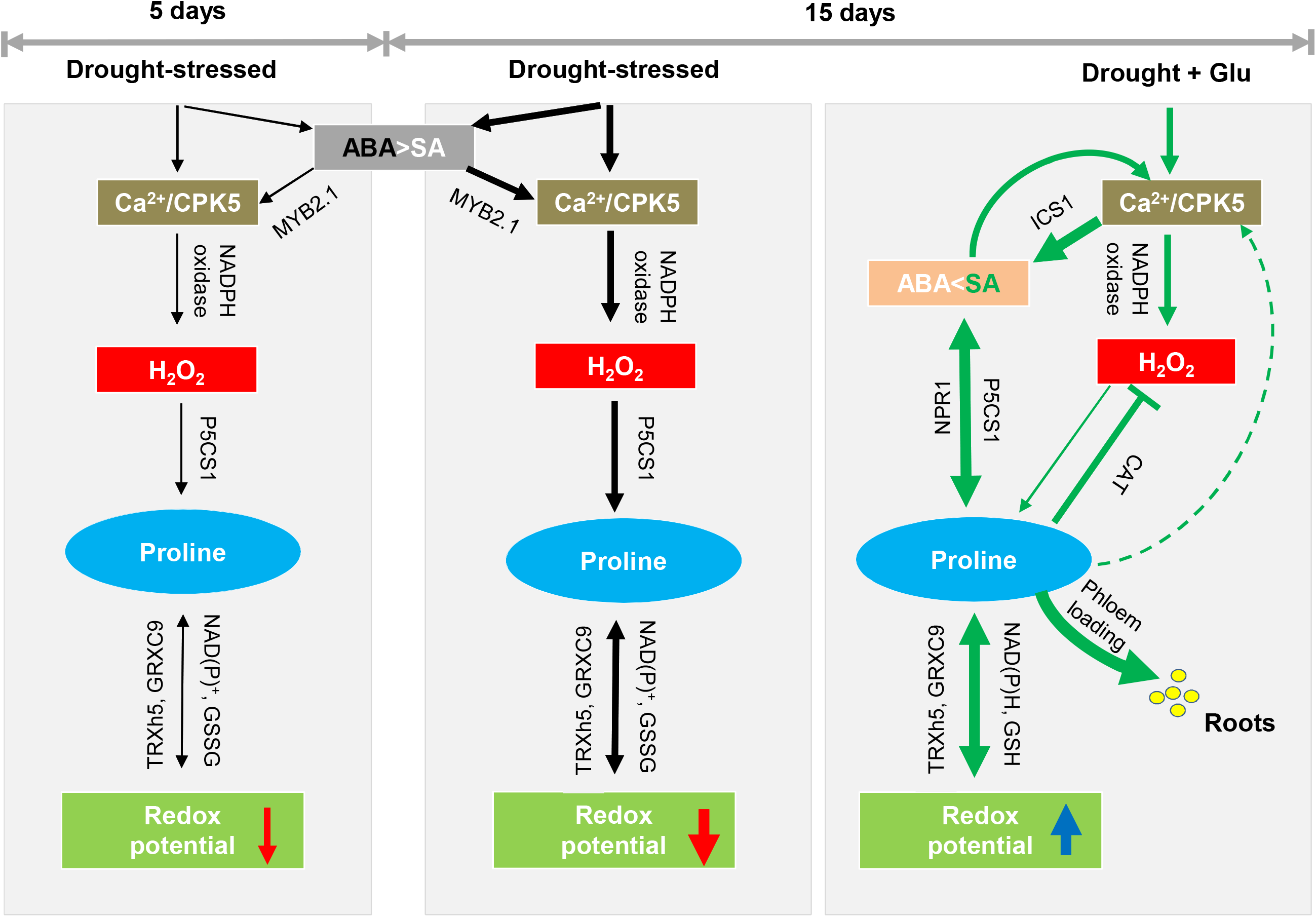
Proposed model for glutamate-mediated hormone antagonism, proline synthesis, and redox modulation under drought and/or glutamate treatment. Black arrows represent the ABA-dependent pathway of response to drought, and green arrows represent the glutamate-mediated SA pathway under drought. Red or blue arrows indicate the decrease or increase of redox potential. The thickness of the arrow expresses the strength of induced or depressed response.

## Supplementary data

**Table S1.** Oligonucleotide primer sequences used for quantitative real-time PCR

**Fig S1.** Changes in antioxidative enzymes activity and catalase (*CAT*) gene expression in the leaves of control or glutamate (Glu)-treated *Brassica napus* under well-watered or drought-stressed conditions. (A) Superoxide dismutase (SOD) and (B) CAT activity and (C) *CAT* gene expression. qRT-PCR was performed in duplicate for each of the three independent biological samples. Values are represented as mean ± SE (n = 3). Different letters on columns indicate significant difference at *P* < 0.05 according to the Duncan’s multiple range test.

**Fig S2.** Proline content in phloem and xylem in control or glutamate (Glu)-treated *Brassica napus* under well-watered or drought-stressed conditions. Proline content in (A) phloem exudates and (B) xylem sap. Values are represented as mean ± SE (n = 3). Different letters on columns indicate significant difference at *P* <0.05 according to the Duncan’s multiple range test.

## Acknowledgment

The present study was supported by the National Research Foundation of South Korea (NRF2017RIA2B4002914).

## References

Asano T, Hayashi N, Kobayashi M, et al. 2012. A rice calcium-dependent protein kinase OsCPK12 oppositely modulates salt-stress tolerance and blast disease resistance. Plant Journal 69, 26–36.

Boudsocq M, Sheen J. 2013. CDPKs in immune and stress signalling. Trends in Plant Science 18, 1.

Chen J, Zhang Y, Wang C, Lü W, Jin JB, Hua, X. 2011. Proline induces calcium-mediated oxidative burst and salicylic acid signaling. Amino Acids 40, 1473–1484.

Chung JS, Zhu JK, Bressan RA, Hasegawa PM, Shi H. 2008. Reactive oxygen species mediate Na+-induced SOS1 mRNA stability in Arabidopsis. Plant Journal 53, 554–565.

Dubiella U, Seybold H, Durian G, Komander E, Lassig R, Witte CP, Schulze WX, Romeis T. 2013. Calcium-dependent protein kinase/NADPH oxidase activation circuit required for defense signal propagation. Proceeding of the National Academy of Sciences of the United States of America 110, 8744–8749.

Elizabeth M, Zhang S. 2000. Calcium-independent activation of salicylic acid–induced protein kinase and a 40-kilodalton protein kinase by hyperosmotic stress. Plant Physiology 122, 1355–1363.

Finiti I, Leyva MO, Vicedo B, Gómez-Pastor R, López-Cruz J, García-Agustín P, Real MD, González-Bosch C. 2014. Hexanoic acid protects tomato plants against Botrytis cinerea by priming defence responses and reducing oxidative stress. Molecular Plant Pathology 15, 550–562.

Herrera-Vásquez A, Carvallo L, Blanco B, Tobar M, Villarroel-Candia E, Vicente-Carbaijosa J. Salinas P, Holuigue L. 2015. Transcriptional Control of Glutaredoxin GRXC9 expression by a salicylic acid-dependent and NPR1-independent pathway in Arabidopsis. Plant Molecular Biology Reporter 33, 624–637.

Hong Z, Lakkineni K, Zhang Z, Verma DPS. 2000. Removal of feedback inhibition of Δ^1^-pyrroline-5-carboxylate synthestase results in increased proline accumulation and protection of plants from osmotic stress. Plant Physiology 122, 1129–1136.

Islam MT, Lee BR, Park SH, La VH, Bae DW, Kim TH. 2017. Cultivar variation in hormonal balance is a significant determinant of disease susceptibility to Xanthomonas campestris pv. campestris in Brassica napus. Frontiers in Plant Science 8, 2121.

Kan CC. Chung TY, Wu HY, Juo YA, Hsieh MH. 2017. Exogenous glutamate rapidly induces the expression of genes involved in metabolism and defense response in rice roots. BMC Genomics, 18, 186.

Kaplan F, Kopka J, Sung DY, Zhao W, Popp M, Porat R, Guy CL. 2007. Transcript and metabolite profiling during cold acclimation of Arabidopsis reveals an intricate relationship of cold-regulated gene expression with modifications in metabolite content. Plant Journal 50, 967–981.

Kim TH, Lee BR, Jung WJ, Kim KY, Avice JC, Qurry A. 2004. De novo protein synthesis in relation to ammonia and proline accumulation in water stressed white clover. Functional Plant Biology 31, 847–855.

Kneeshaw S, Gelinau S, Tada Y, Loake GJ, Spoel SH. 2014. Selective protein denitrosylation activity of thioredoxin-h5 modulate plant immunity. Molecular Cell 56, 153–162.

Knight H, Trewavas AJ, Knight MR. 1996. Cold calcium signaling in Arabidopsis involves tow cellular pools and a change in calcium signature after acclimation. Plant Cell 8, 489–503.

Kotov AA, Kotova LM. 2015. Role of acropetal water transport in regulation of cytokinin levels in stems of pea seedlings. Russian Journal of Plant Physiology 62, 390–400.

La VH, Lee BR, Islam MT, Park SH, Jung HI, Bae DW, Kim TH. 2019. Characterization of salicylic acid-mediated modulation of the drought stress responses: Reactive oxygen species, proline, and redox state in Brassica napus. Environmental and Experimental Botany 157, 1–10.

Lee BR, Li LS, Jung WJ, Jin YL, Avice JC, Qurry A, Kim TH. 2009a. Water deficit-induced oxidative stress and the activation of antioxidant enzymes in white clover leaves. Biologia Plantarum 53, 505–510.

Lee BR, Jin YL, Avice JC, Cliquet JB, Ourry A, Kim TH. 2009b. Increased proline loading to phloem and its effects on nitrogen uptake and assimilation in water-stressed white clover (Trifolium repens). New Phytologist 182, 654–663.

Lee BR, Jin YL, Park SH, Zaman R, Zhang Q, Avice JC, Ourry A, Kim TH. 2015. Genotypic variation in N uptake and assimilation estimated by 15N tracing in water deficit-stressed Brassica napus. Environmental and Experimental Botany 109, 73–79.

Lee BR, Zaman R, Avice JC, Ourry A, Kim TH. 2016. Sulfur use efficiency is a significant determinant of drought stress tolerance in relation to photosynthetic activity in Brassica napus cultivars. Frontiers in Plant Science 7, 459.

Lee BR. Muneer S, Park SH, Zhang Q, Kim TH. 2013. Ammonium-induced proline and sucrose accumulation, and their significance in antioxidative activity and osmotic adjustment. Acta Physiologiae Plantarum 35, 2655–2664.

Liang X, Zhang L, Natarajan SK, Becker DF. 2013. Proline mechanism of stress survival. Antioxidants & Redox Signaling 19, 9.

Livak JK, Schmittgen TD. 2001. Analysis of relative gene expression data using real-time quantitative PCR and the 2^-ΔΔCt^ method. Methods 25, 402–408.

Mezl VA, Knox WE. 1976. Properties and analysis of a stable derivative of pyrroline-5-carboxylic acid for use in metabolic studies. Analytical Biochemistry 74, 430–440.

Miura K, Tada Y. 2014. Regulation of water, salinity, and cold stress responses by salicylic acid. Frontiers in Plant Science 5, 4.

Mou Z, Fan W, Dong X. 2003. Inducers of plant systemic acquired resistance regulate NPR1 function through redox changes. Cell 113, 935–944.

Osakabe Y, Osakabe K, Shinozaki K, Tran LSP. 2014. Response of plants to water stress. Frontiers in Plant Science 5, 86.

Prodhan MY, Munemasa S, Nahar MNEN, Nakamura Y, Murata Y. 2018. Guard cell salicylic acid signaling is integrated into abscisic acid signaling via the Ca2+/CPK-dependent pathway. Plant Physiology 178, 441–450.

Rejeb BK, Lefebvre-De Vos D, Le Disquet I, Leprince AS, Bordenave M, Maldinev R, Jdey A, Abdelly C, Savouré A. 2015. Hydrogen peroxide produced by NADPH oxidases increases proline accumulation during salt or mannitol stress in Arabidopsis thaliana. New Phytologist 208, 1138–1148.

Rejeb KB, Abdelly C, Savouré A. 2014. How reactive oxygen species and proline face stress together. Plant Physiology and Biochemistry 80, 278–284.

Seyfferth C, Tsuda K. 2014. Salicylic acid signal transduction: the initiation of biosynthesis, perception and transcription reprogramming. Frontiers in Plant Science 5, 697.

Sharma S, Villamor JG, Verslues PE. 2011. Essential role of tissue-specific proline synthesis and catabolism in growth and redox balance at low water potential. Plant Physiology 157, 292–304.

Stael S, Kmiecik P, Willems P, Van der Kelen K, Coll NS, Teige M, Breusegem FV. 2015. Plant innate immunity-sunny side up?. Trends in Plant Science 20, 1.

Straus MR, Rietz S, van Themaat E, Bartsch M, Parker JE. 2010. Salicylic acid antagonism of EDS1-driven cell death is important for immune and oxidative stress responses in Arabidopsis. Plant Journal 62, 628–640.

Szabados L, Savouré A. 2010. Proline: a mulitfunctional amino acid. Trends Plant Science 15, 89–97.

Szymanski DB, Liao B, Zielinski RE. 1996. Calmodulin isoforms differentially enhance the binding of cauliflower nuclear proteins and recombinant TGA3 to a region derived from the Arabidopsis Cam-3 promoter. Plant Cell 8, 1069–1077.

Tada Y, Spoel SH, Pajerowska-Mukhtar K, Mou Z, Song J, Wang C, Zuo J, Dong X. 2008. Plant immunity requires conformational charges of NPR1 via S-Nitrosylation and Thioredoxins. Science 321, 952–956.

Tanaka K, Swanson SJ, Gilroy S, Stacey G. 2010. Extracellular nucleotides elicit cytosolic free calcium oscillation in Arabidopsis. Plant Physiology 154, 705–719.

Verslues PE, Kim, YS, Zhu JK. 2007. Altered ABA, proline and hydrogen peroxide in an Arabidopsis glutamate: glyoxylate aminotransferase mutant. Plant Molecular Biology 64, 205–217.

Wang L, Tsuka K, Truman W, Sato M, Nguyen Le V, Katagiri F, Glazebrook J. 2011. CBP60g and SARD1 play partially redundant critical roles in salicylic acid signaling. Plant Journal 67, 1029–1041.

Xia XJ, Zhou YH, Shi K, Zhou J, Foyer CH, Yu JQ. 2015. Interplay between reactive oxygen species and hormones in the control of plant development and stress tolerance. Journal of Experimental Botany 66, 2839–2856.

Zhang Y, Xu S, Ding P, et al. 2010. Control of salicylic acid synthesis and systemic acquired resistance by two members of a plant-specific family of transcription factors. Proceeding of the National Academy of Sciences of the United States of America 107, 18220–18225.

